# Untangling the Role of Pathobionts from Bacteroides Species in Inflammatory Bowel Diseases

**DOI:** 10.1101/2023.10.29.564605

**Authors:** Yue Shan, Joash Lake, Candace M. Cham, Mingming Zhu, Deepinder Kaur, Daina Ringus, Na Fei, Christopher R. Weber, Melanie Spedale, Betty Theriault, Sushila Dalal, David T. Rubin, Aretha Fiebig, Sean Crosson, Karen Lolans, Laurie Comstock, A. Murat Eren, Anindita Basu, Sebastian Pott, Mitchell Sogin, Cambrian Liu, Eugene B. Chang

**Affiliations:** Section of Gastroenterology, Hepatology, and Nutrition, Department of Medicine, The University of Chicago, Chicago, IL 60637 USA; Committee on Immunology, The University of Chicago, Chicago, IL 60637, USA; Committee on Microbiology, The University of Chicago, Chicago, IL 60637, USA; Department of Pathology, The University of Chicago, Chicago, IL 60637 USA; Gnotobiotic Mouse Facility, The Microbiome Center, The University of Chicago, Chicago, IL 60637 USA; Microbiology and Molecular Genetics, Michigan State University, East Lansing, MI 48824 USA; Biosciences Division, Argonne National Laboratory, Argonne, IL 60439 USA; Duchossois Family Institute, Knapp Center for Biomedical Discovery, The University of Chicago, Chicago, IL 60637 USA; Microbiology and Immunology, University of Michigan Medical School, Ann Arbor, MI 48109 USA; Helmholtz Institute for Functional Marine Biodiversity, University of Oldenburg, Oldenburg, Lower Saxony, D-2611 Germany; Pritzker School of Molecular Engineering, The University of Chicago, Chicago, IL 60637 USA; Section of Genetic Medicine, Department of Medicine, The University of Chicago, Chicago, IL 60637 USA; Departments of Pediatrics, Pathology, Human Genetics, and Genetic Medicine, The University of Chicago, Chicago, IL, USA; The Josephine Bay Paul Center for Comparative Molecular Biology and Evolution, Marine Biological Laboratory, 7 MBL Street, Woods Hole, Massachusetts

## Abstract

Inflammatory bowel diseases (IBD) arise from a convergence of underlying genetic susceptibility, environmental factors, and shifts in gut microbiota function and membership. Although the latter may trigger and contribute to IBD, there is little consensus on a specific causative pathogen. In this study, we demonstrate that commensal *Bacteroides fragilis* strains from ulcerative colitis (UC) patients before and during the development of ileal pouchitis engraft and promote colitis in specific pathogen free (SPF) IL-10 deficient (IL-10^-/-^) mice, but not in wild type SPF mice or when mono-associated in germ free mice. The colitis in IL-10^-/-^ mice was also associated with significant alterations in commensal microbiota potentially important for maintaining intestinal and immune homeostasis. UC pouchitis *B. fragilis* also engrafts in DSS-induced colitis in WT SPF mice, indicating a fitness advantage under conditions of mucosal inflammation over other commensals in the gut microbiota. These findings show that gut inflammation promotes the expansion and fitness of UC-derived *Bacteroides* species that is associated with changes in the SPF gut microbiota and may be promote colitis in genetically susceptible hosts.

**Importance:** This study supports the notion that human inflammatory bowel diseases arise from the emergence of indigenous pathobionts in genetically-prone subjects. Colitis-promoting pathobionts are well-suited to establish themselves in the host inflammatory environment and outcompete endogenous microbiota. Once engrafted, the pathobiont can further aggravate inflammation in a genetically-susceptible host. Such complex interplay among several factors creates a vicious pro-inflammatory cycle and promotes disease development. These findings are consistent with our previous clinical observation that *B. fragilis*, an otherwise low-abundance commensal species, expands prior to the development of UC pouchitis. We believe these findings are relevant to the pathogenesis of UC pouchitis and possibly human inflammatory bowel diseases in general, underscoring the role of commensal to pathobiont transitions, rather than classical pathogens, in promoting and exacerbating the onset of human IBD.

## Introduction

The development of inflammatory bowel diseases (IBD) likely involves the convergence of host genetics, environment, and gut microbiota^1^, each necessary, but not sufficient alone to cause disease. Moreover, their relative contributions in the etiopathogenesis of IBD is not clearly defined. Researchers have demonstrated that when compared to healthy individuals, IBD patients exhibit changes in the composition of their gut microbiota, characterized by increase abundance of Proteobacteria and Enterobacteracia and decreased commensal microorganisms^2^. Also, researchers have shown changes in microbial metabolite composition, with IBD patients showing increased microbial-associated metabolites such as sphingolipids and alcohols, while healthy individuals exhibited greater indole metabolites that have known protective properties in murine colitis^3,4^. However, without insight into who will develop the disease and when, it is impossible to determine whether these changes in the gut microbiota are cause or consequence of IBD.

Researchers and medical practitioners have been hindered in studying IBD due to a lack of understanding of causal mechanisms and biological indicators that may be present in genetically-susceptible individuals. To this end, we have studied ulcerative colitis (UC) patients who have undergone an ileal pouch-anal anastomosis (IPAA) to determine the events that occur prior to the onset of IBD^5,6^. Nearly half of UC-IPAA patients develop another form of inflammatory disease of the ileal pouch called pouchitis within two years of pouch functionalization (defined as when the pouch is exposed to the fecal stream). Researchers have long speculated a microbial basis for IBD^7^, arising from compromises in the commensal gut microbial population and/or the emergence of pathobionts that are innocuous in one context, but promote inflammation under certain circumstances^8^. Previously, we employed metagenomic sequencing to assess the changes in membership in UC pouchitis patients^6^. Metagenomic analysis showed that in this cohort of IPAA patients, several *Bacteroides* species expanded prior to the onset of pouchitis, with the most frequent being non-enterotoxigenic *Bacteroides fragilis*^6^. As such, this makes *Bacteroides fragilis* a putative pathobiont, where in the gut of a healthy individual it does not promote disease, however its expansion in an UC-IPAA patient may cause it to disrupt intestinal homeostasis, resulting in pouchitis.

*Bacteroides fragilis* is a gut commensal present in most healthy individuals at very low abundance (<0.5%)^9,10^. It has been previously shown that enterotoxigenic strains of *B. fragilis* secrete Bacteroides fragilis toxin (Bft), which is associated with the onset of colorectal cancer and colitis^11–13^. However, non-enterotoxigenic strains of *B. fragilis* have been shown to have immunomodulatory properties, promoting the induction of IL-10^+^ CD4^+^ T Cells and ameliorating murine colitis^14,15^. The immunomodulatory properties were mediated by Capsular Polysaccharide A (PSA) interacting with TLR2^14,16^. In IBD patients (both Crohn’s Disease and ulcerative colitis), researchers have shown that *B. fragilis* can become highly represented in the gastrointestinal tract compared to healthy individuals^2^ and, as shown by others, highly coated with IgA, suggesting that it may also have immunomodulatory properties beyond the induction of IL-10^+^ CD4^+^ T cells^17^. Human-isolated strains of non-enterotoxigenic *B. fragilis* ZY-312 have also been shown to polarize bone-marrow-derived macrophages to a pro-inflammatory M1 phenotype, suggesting that these commensals may have pro-inflammatory properties as well^18,19^. The differing responses to non-enterotoxigenic *B. fragilis* highlight the fact that not all non-enterotoxigenic strains of *B. fragilis* modulate the immune system the same way.

In the current study, we cultivate non-enterotoxigenic *B. fragilis* strains from UC pouchitis patients and examined the role of *B. fragilis* in multiple experimental colitis models. These findings suggest that non-enterotoxigenic *B. fragilis* strains isolated from UC pouch patients have the potential to initiate and contribute to the development of immune-mediated colitis in genetically-susceptible hosts under the right conditions. This observation makes us rethink the traditional paradigm that complex immune disorders like IBD are caused by classical pathogens. Rather, these diseases can be initiated when certain commensals, given the right circumstance and opportunity, establish a new stable and resilient population in genetically-susceptible hosts and promote the severity and chronicity of inflammation.

## Results

### Isolation of non-enterotoxigenic *B. fragilis* from ulcerative colitis pouchitis patients

In our previous prospective study of IPAA patients, we collected samples of pouch microbiota every four months over a two year period^6^. Significant shifts in the pouch microbial communities were noted in patients who developed pouchitis, including marked expansion of *Bacteroides spp such as B. fragilis* in several subjects prior to and during inflammation. Because *B. fragilis* is typically a low-abundance species found among healthy humans, we cultivated *B. fragilis* strains from UC IPAA patients using Bacteroides Bile Esculin (BBE) plates and PCR screening. For each patient sample, multiple independent colonies of *B. fragilis* were cultivated, but whole-genome sequencing revealed that only one strain could be detected in individual pouchitis patients, and none of the isolated *B. fragilis* were enterotoxigenic pathogens (data not shown).

To evaluate the physiological impact of *B. fragilis* on pouchitis patients, we sought to test the impact of these strains in an animal model^20^. We started by mono-associating these strains in germ-free wild-type (WT) C57Bl/6 mice. The studies were performed in WT and *interleukin 10* knockout (IL-10^-/-^) mice, both on the C57Bl/6 background. IL-10 is an important cytokine produced by regulatory T cells and is known to have anti-inflammatory effects. Polymorphism or loss of function in the *IL10* gene is associated with onset of monogenic human IBD early in life^21–24^. IL-10^-/-^ mice are also prone to the development of spontaneous colitis, but only in the presence of specific pathogen free (SPF) gut microbiota, and germ-free IL-10^-/-^ mice do not develop spontaneous colitis^25^. We mono-associated patient-derived *B. fragilis* isolates into germ-free WT and IL-10^-/-^ mice and found engraftment of *B. fragilis* did not cause any colitis in either line, despite stable colonization (Figure 1A, B).

**Figure 1.**
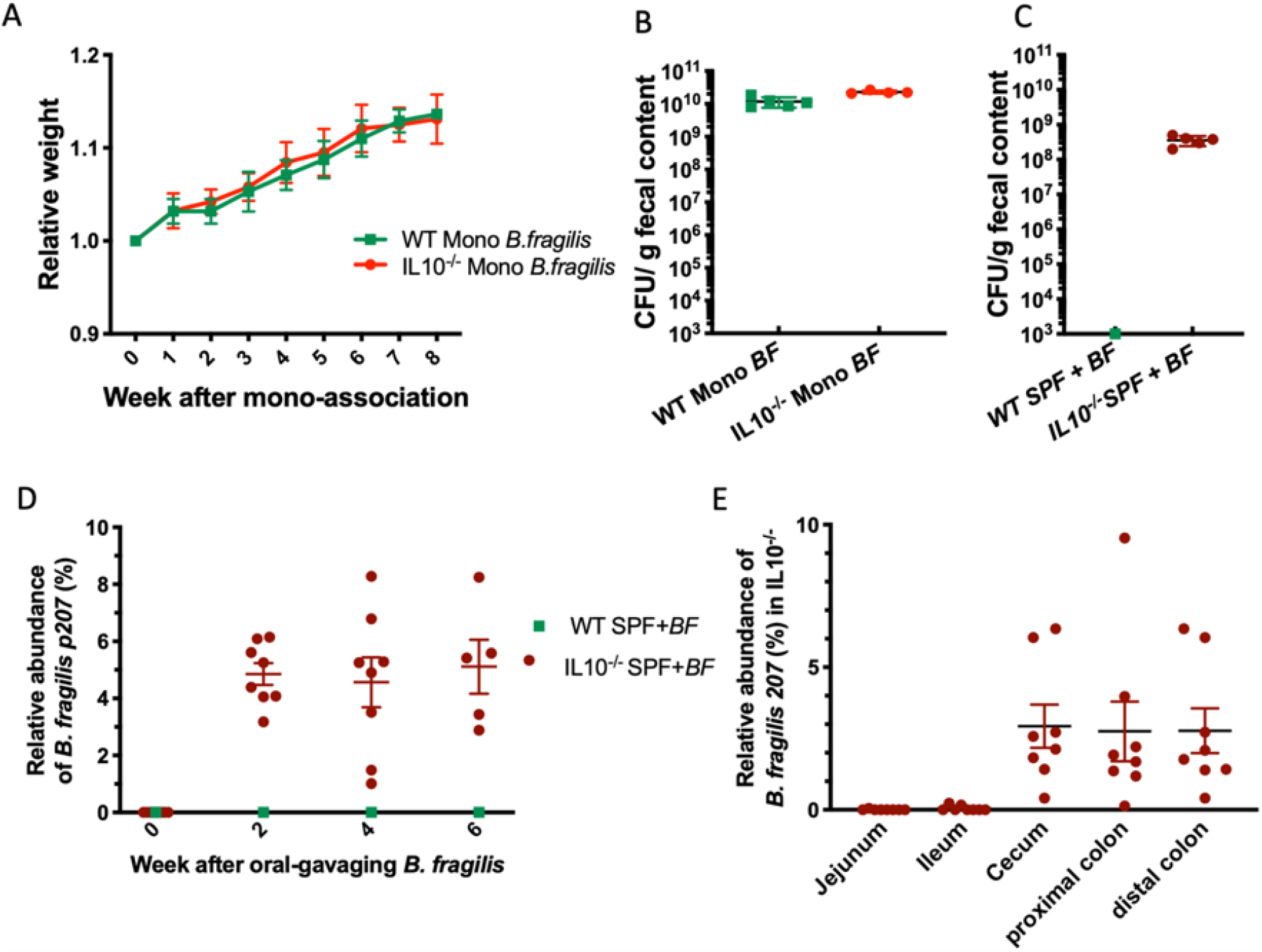
*B. fragilis* engrafted into IL-10^-/-^, but not WT, mice with an SPF consortium. (A) Monoassociation of *B. fragilis* did not cause weight loss in either WT or IL-10^-/-^ mice. (B) CFU of *B. fragilis* in fecal content of mono-associated WT and IL-10^-/-^ mice. (C) CFU of *B. fragilis* in fecal content of SPF WT and IL-10^-/-^ mice. (D) The relative abundance of *B. fragilis* measured by qPCR in fecal content of SPF WT and IL-10^-/-^ mice. (E) The relative abundance of *B. fragilis* measured by qPCR in luminal content of SPF IL-10^-/-^ mice.

We reasoned that the mono-association of *B. fragilis* may not have caused colitis because commensal gut microbiota is required for maturation of the immune cell populations in the gut^20^. We therefore sought to evaluate the impact of *B. fragilis* in the presence of a SPF microbiota (Figure S1). To minimize any cage effects, we conventionalized both WT and IL-10^-/-^ germ-free mice with the same SPF cecal consortia. After the establishment of the complex microbiota, we administered *B. fragilis* by oral gavage to both groups. We followed the groups for 6 weeks by plating fecal samples on *B. fragilis* selective plates and screened by using specific PCR primers. We found that *B. fragilis* failed to engraft in WT SPF mice (Figure 1C), probably as a result of the strong priority effect of the established SPF murine microbiota.

In contrast to conventionalized WT SPF mice, we found that *B. fragilis* engrafted into the SPF microbiota of SPF conventionalized IL-10^-/-^ mice (Figure 1C). The engraftment of *B. fragilis* remained stable over time and evenly distributed in the cecum and colon of the host (Figures 1D, E), reaching relative abundances of ∼5% in the fecal and luminal microbiota in IL-10^-/-^ mice at levels even higher than typically found in healthy human subjects (<0.5%). This finding could be due to the pro-inflammatory environment of the SPF IL-10^-/-^ host gut ecosystem that is favorable for the human *B. fragilis* isolates, but unfavorable for indigenous commensal murine microbiota.

### *B. fragilis* aggravates the severity of host inflammation and spontaneous colitis in IL-10^-/-^ mice

IL-10^-/-^ mice are susceptible to the development of chronic inflammation in the presence of a complex gut microbiota, and the reported prevalence of spontaneous colitis varies among facilities due to differences in gut microbiota composition. In our cohort, we observed a ∼25% incidence of rectal prolapse, an indicator of severe inflammation of the colon. In the group given *B. fragilis*, the rectal prolapse incidence rate increased to ∼75% (Figures 2A, B), suggesting that *B. fragilis* augmented, rather than protected against, the onset of spontaneous colitis. We observed a significant increase in levels of lipocalin-2 (Lcn2), a non-invasive fecal biomarker for colitis, and TNFα, a signature cytokine for inflammation (Figures 2C-E) in IL-10^-/-^ mice colonized with *B. fragilis*. Furthermore, colon histology showed a trend of increased severity (Figures 2F, S2). Combining these data, we concluded that colonization of *B. fragilis p207* into a complex gut microbiota aggravates inflammation and colitis in genetically-susceptible mice.

**Figure 2.**
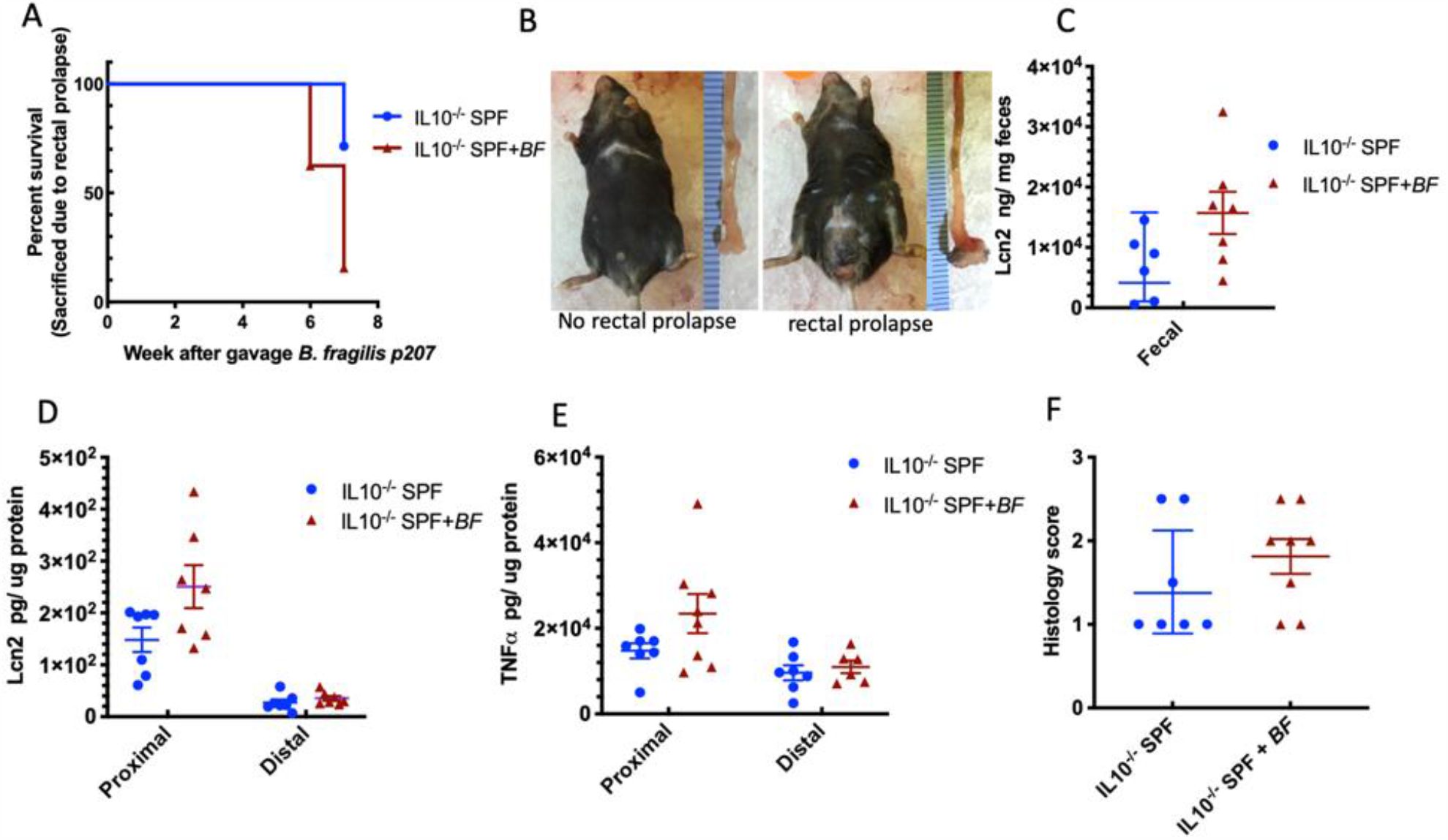
Engraftment of *B. fragilis* aggravates inflammation in IL-10^-/-^ SPF mice. (A) Percentage of mice with spontaneous rectal prolapse. (B) Representative images of mice with or without rectal prolapse. (C) The protein level of lipocalin 2 (Lcn2) in fecal content as measured by ELISA. (D) The protein level of Lcn2 in colonic mucosa as measured by ELISA. (E) The protein level of TNFα in colonic mucosa measured by ELISA. (F) Histology score of colon tissue.

### Engraftment of *B. fragilis* impacts the composition of the SPF gut microbiota

To understand the overall change of gut microbiota after the engraftment of *B. fragilis*, we performed 16S rRNA gene amplicon sequencing of fecal samples in SPF conventionalized IL-10^-/-^ mice with or without *B. fragilis*. Principal coordinate analysis (PCoA) plot using weighted UniFrac distances demonstrated that the two groups formed separate clusters (Figure 3A), even though these groups were indistinguishable before introduction of *B. fragilis* (Figure 3B, week 0). We also compared the relative abundance of each microbial family between the two groups. Interestingly, introducing *B. fragilis* reduced the relative abundance of *Bacteroidaceae* and *Lactobacillaceae* families while increasing the relative abundance of *Muribaculaceae* (formerly known as *S24-7*) family (Figure S3).

**Figure 3.**
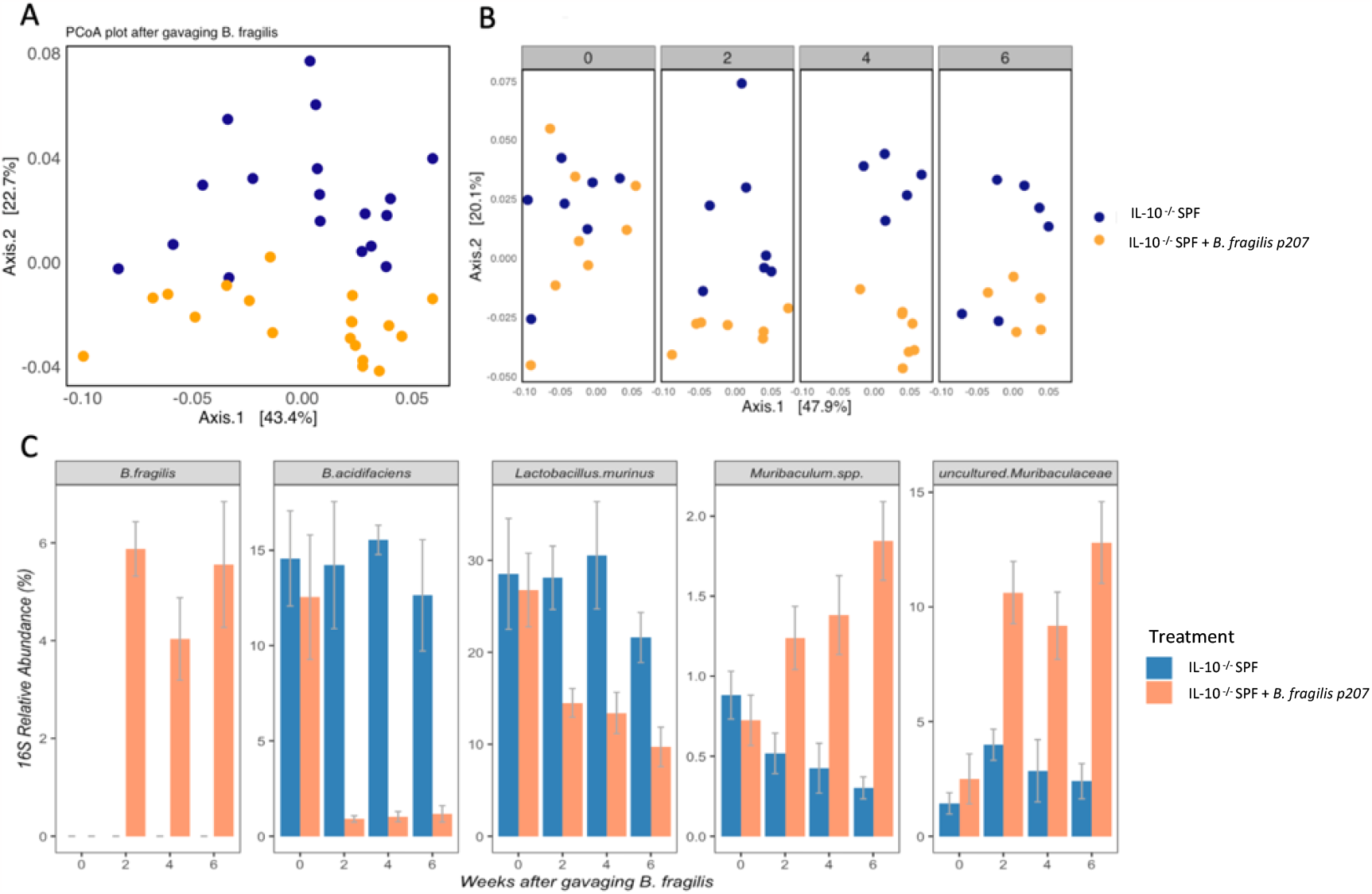
*B. fragilis* dramatically changes the composition of gut microbiota in IL-10^-/-^ SPF mice. (A) PCoA analysis of 16S amplicon sequencing results using the Weighted-UniFrac method. (B) PCoA analysis of week 0, 2, 4, 6 after gavaging *B. fragilis*. (C) The relative abundance of OTUs with significant changes after engraftment of *B. fragilis*.

To overcome the low resolution of 16S amplicon sequencing in identifying taxonomy, we conducted PacBio Single Molecule, Real-Time (SMRT) sequencing, and obtained the full 16S sequence for the OTUs with significant changes upon *B. fragilis* engraftment (Figure 4C). Interestingly, engraftment by *B. fragilis* nearly eliminates an OTU identified as *Bacteroides acidifaciens*, which is highly abundant in the IL-10^-/-^ SPF only group. *B. acidifaciens* is an endogenous murine gut species that was reported to protect mice from diabetes and obesity ^26^. Elimination of *B. acidifaciens* may be due to the direct interspecies competition between the two *Bacteroidetes*^27^. In addition, we observed significant increases in two OTUs that belong to the *Muribaculaceae* family (formerly *S24-7*) and a decrease in one OTU identified as *Lactobacillus murinus* (Figure 3C). The *Muribaculaceae* family has been reported and characterized recently, but its role in gut microbiota and its interaction with the host remains unknown^28^. *Lactobacillus murinus* is commonly used as a probiotic and is believed to have anti-inflammatory effects. It remains unclear whether the change in microbiota structure is a cause or consequence of the host inflammation. This would need to be addressed by isolating and cultivating these OTUs and examining their interaction with the host immune system. From this, we conclude that the microbiota composition and the host physiology undergo significant changes after the engraftment by *B. fragilis*.

**Figure 4.**
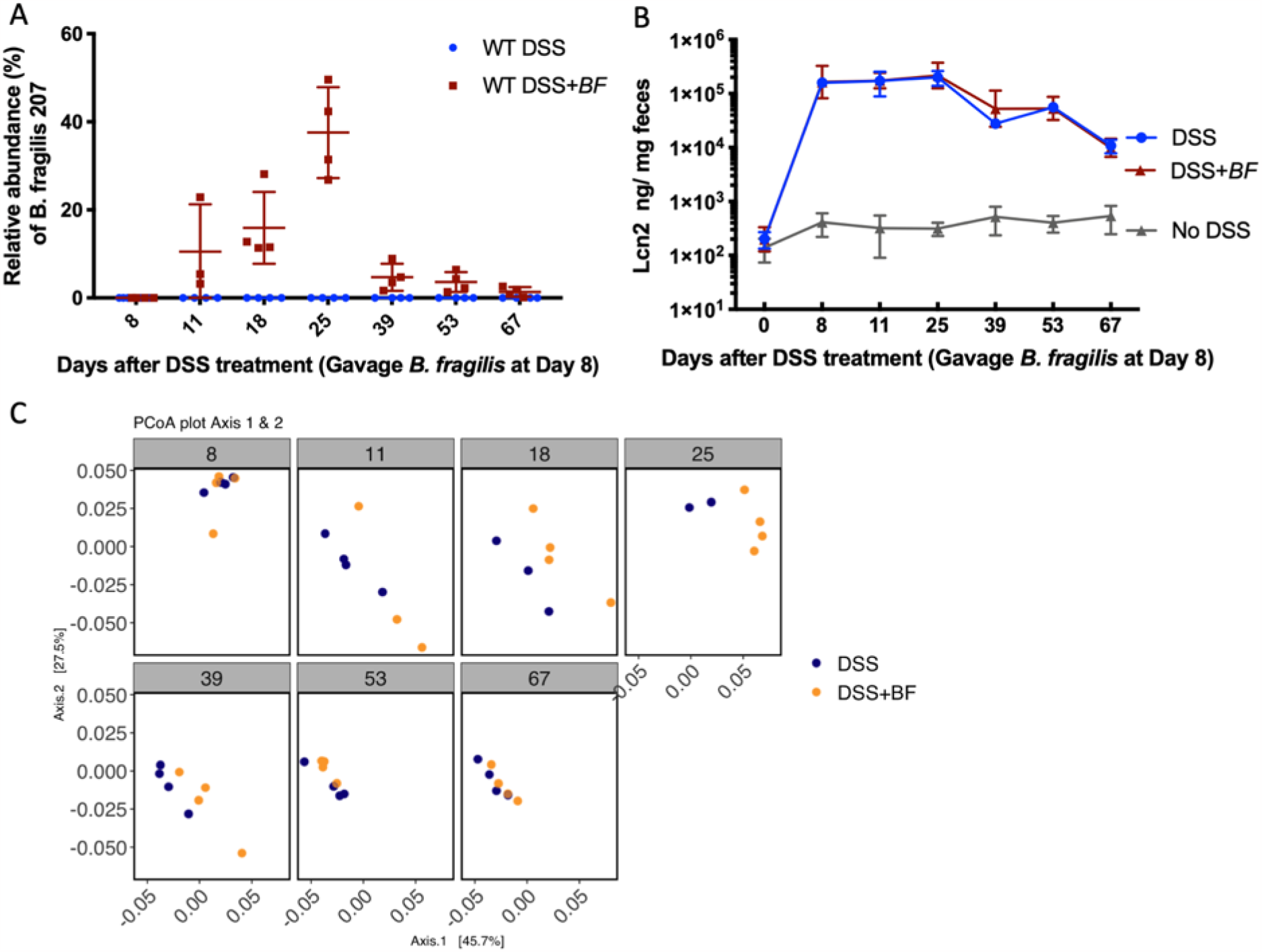
*B. fragilis* engrafts into WT SPF mice after DSS-induced colitis. (A) The relative abundance of *B. fragilis* in WT SPF mice with DSS-induced colitis. (B) The protein level of Lcn2 in fecal content as measured by ELISA. (C) PCoA analysis of 16S amplicon sequencing results of fecal samples from DSS induced colitis in WT SPF mice with or without *B. fragilis* using the Weighted-UniFrac method.

### *B. fragilis* blooms in WT SPF mice after DSS-induced colitis

The different outcomes of *B. fragilis* engraftment between WT SPF and IL-10^-/-^ SPF is worth noting, as this suggests that *B. fragilis* may preferably colonize hosts in an inflammatory environment. To test the hypothesis that the inflammatory environment in IL-10^-/-^ SPF promoted the engraftment of *B. fragilis*, we chemically-induced acute colitis by administrating dextran sodium sulfate (DSS) in the drinking water of WT SPF mice. In DSS-treated WT mice, we saw increased LCN2 levels compared to non-DSS-treated controls, confirming that inflammation was induced in the mice (Figure 4B). During the peak of colitis at day 8 post-DSS treatment (Fig. S4), we administered patient isolate *B. fragilis p207* and followed its relative abundance in fecal samples over time. We found that following DSS-induced colitis, *B. fragilis* engrafted into SPF-conventionalized WT mice. The relative abundance of *B. fragilis* reached up to ∼40% during acute inflammation (Figure 4A). Moreover, as the hosts recovered from acute colitis, the relative abundance of *B. fragilis* also decreased to less than 2%. Similar to SPF conventionalized IL-10^-/-^ mice, although not as dramatic, we saw elevated Lcn2 and TNFα in DSS-treated WT SPF-conventionalized mice with *B. fragilis* at 28 days post DSS-treatment compared to those without *B. fragilis* (Figure S5B, C). When the mice were sacrificed on day 28 post-DSS treatment, the body weights were similar between DSS-treated and none-DSS treated mice (Figure S5A). These data suggest that under intestinal inflammation, *B. fragilis* expansion in the gut has the potential to exacerbate inflammation leading to elevated cytokine levels 28 days following DSS treatment. Finally, we also conducted 16S rRNA gene amplicon sequencing of the fecal samples. PCoA analysis demonstrated a transient difference in the composition of the gut microbiota between the two groups during active inflammation (Figure 4C). Together, these data show that *B. fragilis* has a fitness advantage in an inflammatory environment of DSS-treated WT mice and can expand in the presence of endogenous microbiota that may not be as fit in a colitic environment (compromising its priority effect). The observation supports our hypothesis that *B. fragilis* preferably engrafts host in an inflammatory environment and the engraftment of *B. fragilis* worsens the inflammation of genetically-susceptible hosts.

## Discussion

Inflammatory bowel diseases are multifaceted disorders that rely on the host’s genetic susceptibility, diet, environment, and gut microbiota^1^. While each facet by itself is insufficient to promote disease, the relative contribution of each is still unknown. To understand how the gut microbiota changes prior to the onset of the disease, we studied UC pouchitis patients^5,6^. These patients previously had severe ulcerative colitis (UC). To promote remission, UC patients underwent a total colectomy followed by an ileal pouch-anal anastomosis, in theory curing the disease. Perhaps due to their genetic predisposition and gut microbiota, about half of these patients develop pouchitis within two years of their treatment. Our group has shown prior to the clinical manifestation of the disease, there is an expansion of several *Bacteroides* species, including non-enterotoxigenic *Bacteroides fragilis*^6^. As such, we set out to understand the contribution of non-enterotoxigenic *B. fragilis* in inflammatory bowel diseases.

Based on our findings, *B. fragilis* is likely a pathobiont. In healthy individuals, it may lie at low abundance, but in inflamed environments, it can bloom and flourish in the microbial community (Figure 1C-E). The increased abundance of *B. fragilis* changes the gut microbiota composition, promoting dysbiosis (Figure 3A-C). We showed that in an inflammatory environment, such as DSS-induced colitis and in IL-10^-/-^ mice, Pouchitis-associated pathobiont (PAP) *Bacteroides fragilis* can engraft into the gut microbiota. When administered by mono-association, *B. fragilis* does not cause overt colitis in IL-10^-/-^ and WT mice. However, when given to IL-10^-/-^ mice in the presence of an SPF cecal consortium, we observed an increased incidence of rectal prolapse in mice with the same starting microbiota. This supports the idea that PAP *B. fragilis* cannot cause overt colitis by itself, but instead changes the gut microbiota leading to a dysbiosis and eventual colitis in genetically-susceptible hosts (Figure. 1A-B, 3A-B). Its engraftment and expansion in the IL-10^-/-^ mice prior to rectal prolapse and its bloom in pouchitis patients prior to the onset of disease both suggest that *B. fragilis* modulates the gut microbiota and reinforces the pathways that lead to clinical manifestations of IBD (Figure 2, 3C). Engraftment of PAP *B. fragilis* in IL-10^-/-^ was associated with increased TNFα, suggesting that its presence promoted increased inflammation compared to IL-10^-/-^ control mice.

Non-enterotoxigenic strains of *Bacteroides fragilis* are commonly characterized as symbionts due to their ability to reduce the severity of inflammation, protect against colorectal cancer, and prevent the onset of experimental autoimmune encephalomyelitis^14–16,29^. This antiinflammatory property is mediated by PSA interacting with TLR2, reducing the inflammatory properties of diverse immune cell subsets, including the induction of IL-10^+^ CD4^+^ T cells both directly and through plasmacytoid dendritic cell interactions with CD4^+^ T cells^14,30^. However, others have shown that in inflammatory bowel diseases, *B. fragilis* strains are among the commensals that are highly coated by IgA, suggesting that *B. fragilis* may have other immunomodulatory properties^17^. Researchers have also shown that in IBD patients, non-enterotoxigenic strains of *B. fragilis* have the promoter in the off orientation, upstream of PSA, suggesting that PSA is lowly or not expressed during IBD^31^. In addition, not all non-enterotoxigenic strains ameliorate DSS-induced colitis and some strains even skew macrophages to take on a pro-inflammatory M1-phenotype^18,32^. Furthermore, capsular polysaccharides are distinct between strains of *Bacteroides fragilis*^33^. We have shown that between *B. fragilis* strains, the genetic composition of PAP *B. fragilis* p207 and strain 9343, are genetically-distinct, suggesting that they may have distinct immunomodulatory properties (Figure S6). As such, this further underscores the need to understand strain differences between the non-enterotoxigenic strains of *B. fragilis* to better understand how these commensals influence the immune system in IBD patients.

In our hands, *B. fragilis* could not engraft in the WT mice with a complex gut microbiota (Figure 1C-D). However, when gut inflammation is present, whether through DSS-induced colitis or genetic deletion of *IL10, B. fragilis* can break colonization resistance and engraft into the murine gut microbiota. This property is likely why *B. fragilis* is extremely abundant in IBD patients, as it can thrive better under inflammatory conditions than other microbes. In the IL-10^-/-^ mice, we also showed that the microbiota becomes quite distinct by 6 weeks post-*B. fragilis* engraftment. However, in DSS-treated wildtype mice with *B. fragilis*, our data show that the gut microbiota composition becomes returns to nearly normal by 67 days post *B. fragilis* administration, when inflammation has resolved and the mice have completely recovered from DSS-induced colitis. Thus, under inflammatory conditions, this putative pathobiont can outcompete different microbes in the inflamed intestinal tract and is likely a part of the reason why so many immunomodulatory microbial metabolites are altered in patients with IBD. This supports the notion that dysbiosis gradually occurs prior to the clinical manifestation of IBD.

Our findings implicate non-enterotoxigenic *B. fragilis* as a pathobiont. It may be less fit under non-inflamed conditions and less able to compete with SPF microbiota. However, in an inflamed environment, it can outcompete other microbes, such as *Lactobacillus* species, reinforcing a dysbiosis that often ensues in individuals with IBD. Future studies will be needed to characterize what enables *B. fragilis* to engraft an inflamed environment. Its expansion prior to the onset of the disease should be further investigated in humans to assess whether it can serve as a clinical indicator in individuals that may go on to develop IBD.

### Limitations

Multiple factors, including diet, environment, changes in the gut microbiota, and genetic susceptibility have been associated with Inflammatory Bowel Diseases. In our study, we show that *B. fragilis* can engraft into environments with increased inflammation, such as the IL-10^-/-^ mice and DSS-treated wild-type mice. In this inflammatory environment, based on long-read 16S rRNA sequencing data, this commensal changes the gut microbiota composition. However, how immunomodulatory metabolites change due to the expansion of *B. fragilis* would need to be evaluated to better understand *how B*. fragilis influences intestinal inflammation. This study can be continued by evaluating the changes in the immune cell populations of SPF + *B. fragilis IL-10*^*-/-*^ or DSS-treated WT mice compared to the SPF-only counterparts.

## Figures and Figure legend

**Figure S1.**
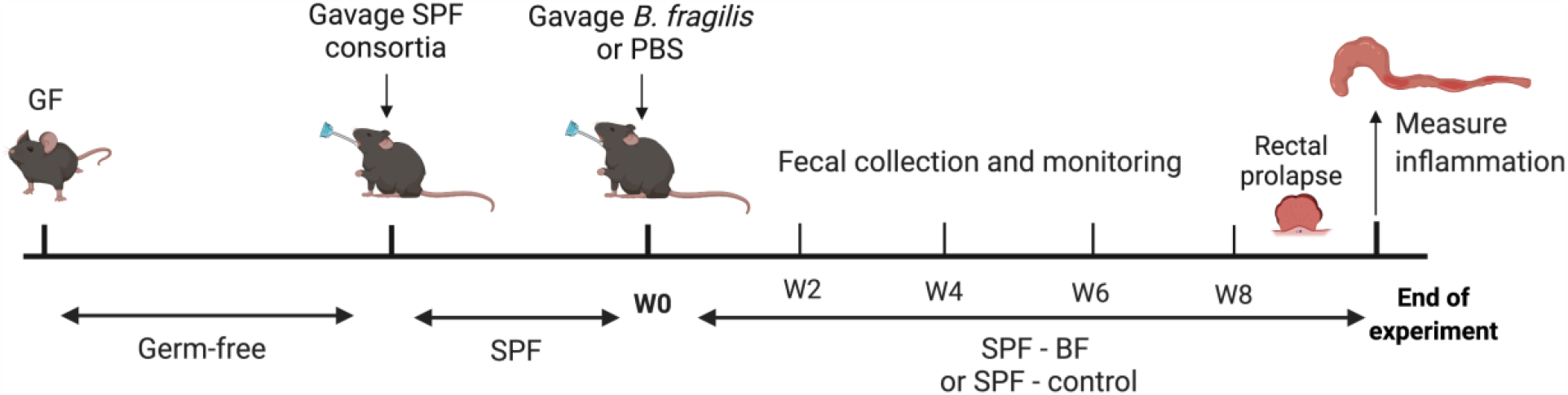
Scheme of conventionalization of WT or IL-10^-/-^ mice with SPF consortia followed by *B. fragilis*. Seven-week germ-free mice were conventionalized with SPF consortia. After stabilization of microbiota for 2 weeks, mice were gavaged with either *B. fragilis* or PBS (control). Mice were monitored and fecal samples were collected weekly for 9 weeks. The onset of rectal prolapse were monitored during the experiments. At the end of experiment, mice were sacrificed and colon samples were collected for analysis.

**Figure S2.**
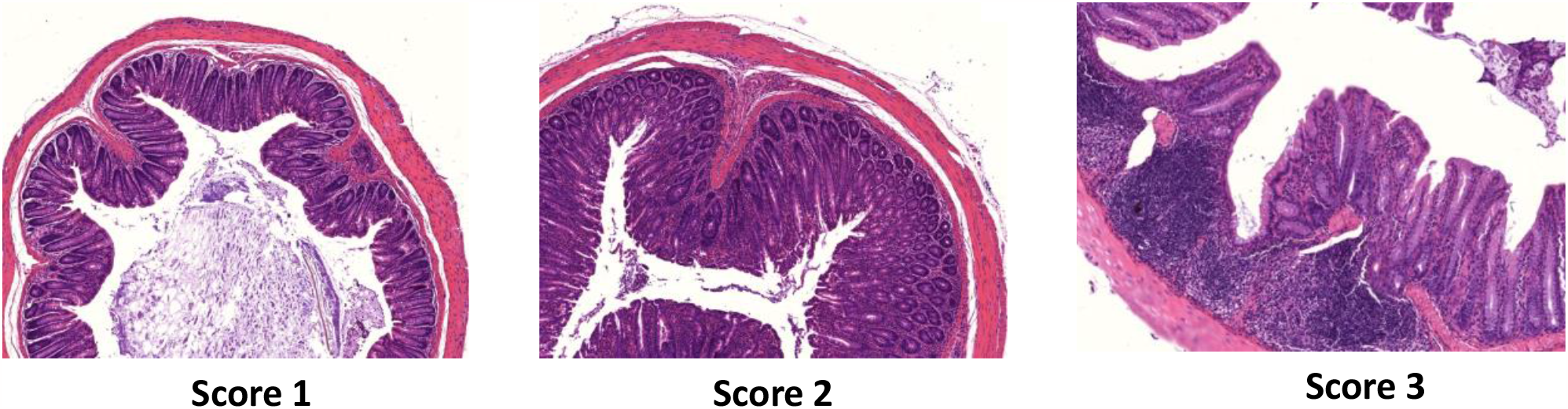
Example of histology score. Cross-section intestinal segments were fixed in 4% formaldehyde and embedded in paraffin followed by H&E staining. Histological grading of intestinal inflammation was performed by a pathologist blinded to the treatment conditions using a validated scoring system previously described: no inflammation was scored as 0; modest numbers of infiltrating cells in the lamina propria was scored as 1; infiltration of mononuclear cells leading to separation of crypts and mild mucosal hyperplasia was scored as 2; massive infiltration with inflammatory cells accompanied by disrupted mucosal architecture, loss of goblet cells, and marked mucosal hyperplasia was scored as 3.

**Figure S3.**
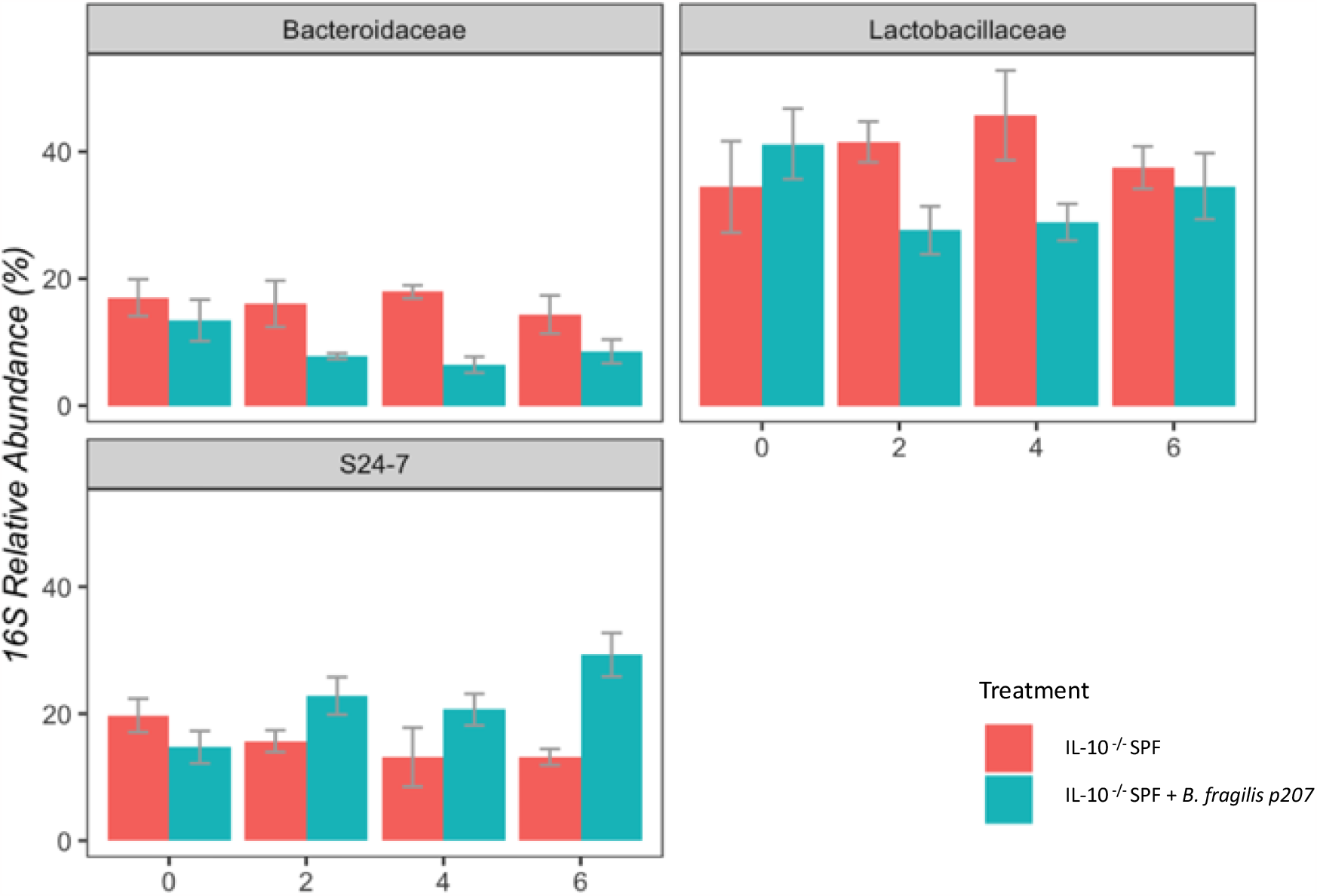
Significant changes of microbial taxa after engraftment of *B. fragilis* in IL-10^-/-^ mice at the family level. The result is based on16S amplicon sequencing.

**Figure S4.**
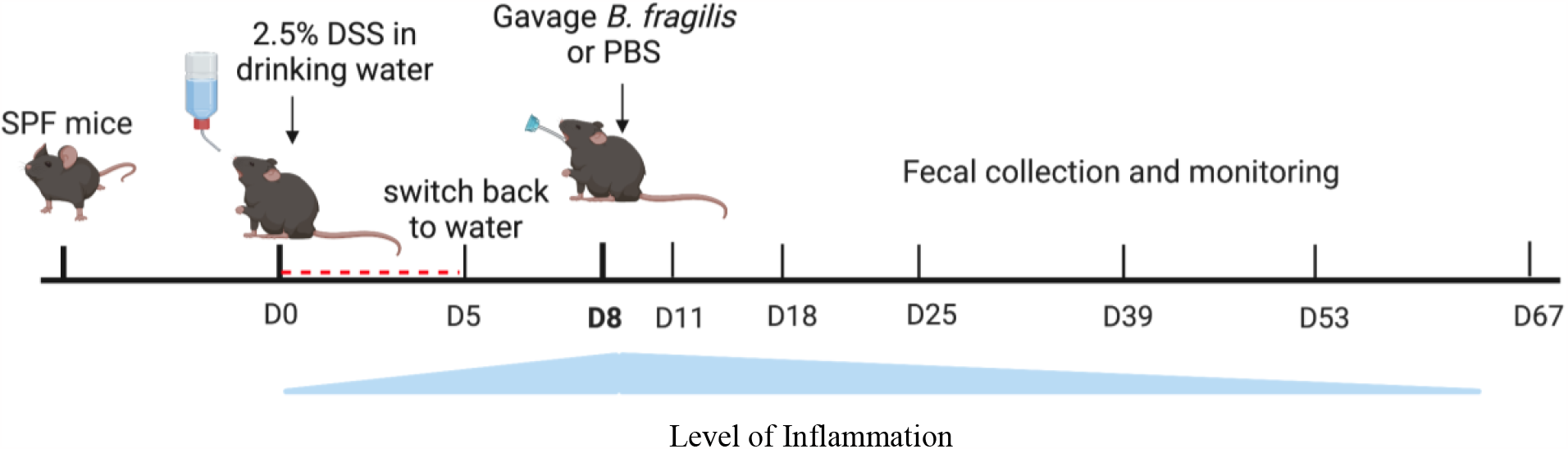
Scheme of DSS-induced colitis model. SPF mice were given 2.5% DSS in drinking water for 5 days (Day 0 to Day 5). Mice were then switch back to water. After 3 days (Day 8), mice were gavaged with either *B. fragilis* or PBS (control). Fecal samples were collected and mice were monitored during the self-recovery.

**Figure S5.**
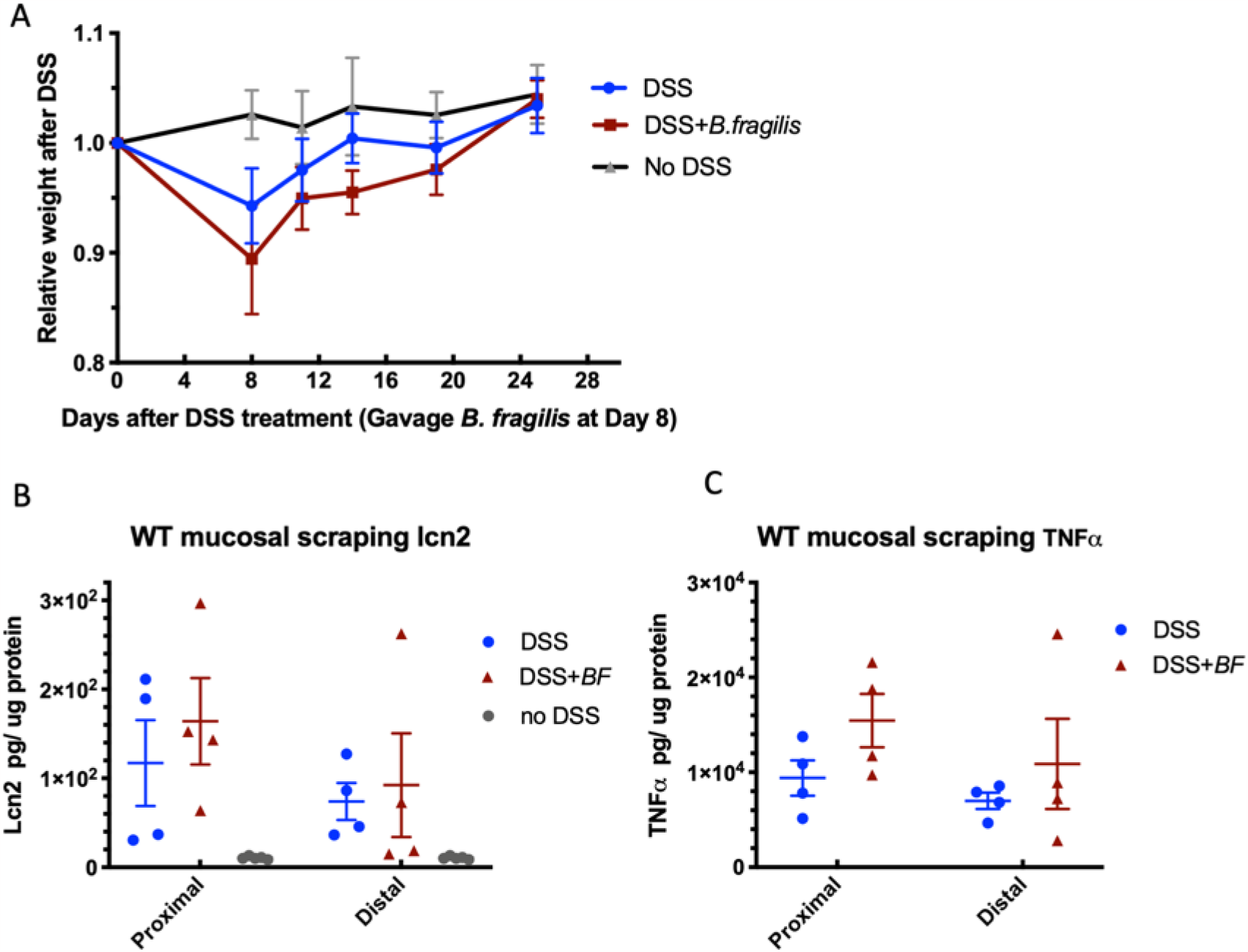
Inflammation levels are augmented in DSS-treated WT SPF + *B. frag* mice. (A) Relative weight changes after DSS treatment. (B) The protein level of lipocalin 2 (Lcn2) in colonic mucosa as measured by ELISA. (C) The protein level of TNFα in colonic mucosa measured by ELISA.

**Figure S6.**
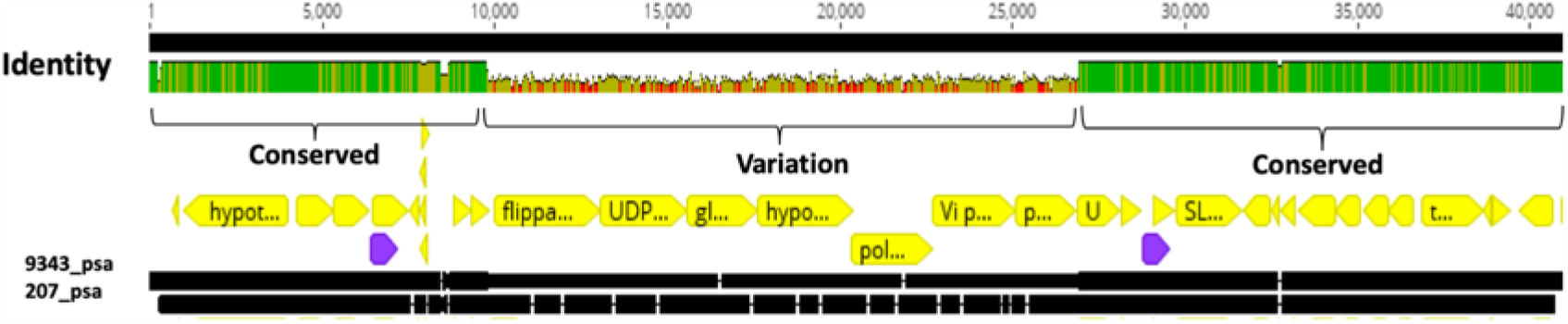
The PSA promoter region is conserved among different strains whereas the structural genes are highly diverse.

## Acknowledgments

We would like to acknowledge the dedicated staff of the Gnotobiotic Research Animal Facility at the University of Chicago for their support in maintaining the germ-free and conventionalized mice used in these studies. We thank The University of Chicago Human Tissue Resource Center (RRID:SCR_019199). This project was supported by the following grants from the National Institute of Diabetes and Digestive and Kidney Diseases: RC2DK122394, R01DK47722, and R01DK113788; and the Center for Interdisciplinary Study of Inflammatory Intestinal Diseases (P30 DK042086). The work was also supported by the 1F31DK136333-01 awarded by the NIH NIDDK, T32 DK07074 awarded by the University of Chicago Section of Gastroenterology, **T**32 AI7090-44 the University of Chicago Committee on Immunology Program, and University of Chicago Initiative for Maximizing Student Development (IMSD) 5R25GM109439.

## Materials and Methods

### Bacterial Isolation and Cultivation

*B. fragilis* cultivars were routinely grown in supplemented <u>B</u>rain <u>H</u>eart Infusion (BHI-S) medium or agar at 37°C in an anaerobic chamber. <u>B</u>acteroides <u>B</u>ile <u>E</u>sculin (BBE) agar was used to selectively grow and quantify *B. fragilis* from fecal and cecal samples of mice.

### Animal

All animal studies have been approved by the Institutional Animal Care and Use Committee at the University of Chicago. All mice used for these studies were on a C57Bl/6J genetic background (WT and IL-10^-/-^). Germ-free (GF) and Specific Pathogen Free (SPF) mice used were bred in the University of Chicago Animal Resource Center.

### Mono- and multiplex-association

Mono- and multiplex association were performed by orally gavaging germ-free animals with bacterial culture or microbiota from mice donors. To make the *B. fragilis* gavaging solution, *B. fragilis* were cultured in BHI-S to late exponential phase (5 X 10^8^ CFU/ml). The bacteria were spun down and resuspended in the same volume of sterile PBS+10% glycerol. For the SPF conventionalization solution, cecal contents were collected anaerobically from WT male mice from our SPF facility at the age of 7-9 weeks. Samples from 14 mice from 7 cages were pooled and suspended in sterile PBS+10% glycerol (100 mg cecal content per 1 ml solution).

For mono-association, GF male mice at ∼9-week of age were gavaged with 100 ul *B. fragilis* solution. For multiplex-association, GF male mice at ∼7-week of age were first gavaged with 100 ul SPF conventionalization solution prepared from gender- and age-matched SPF cecal content. After two weeks, the mice (∼9-week age) received either 100 ul *B. fragilis* gavaging solution or 100 ul control solution (PBS+10% glycerol). For DSS-treatment groups, WT GF male mice at ∼5-week of age were first conventionalized with SPF solution. DSS (2.5% in drinking water) were given for 5 days at the age of 8 weeks. At the end of day 5 of the DSS treatment, mice were switched back to normal drinking water. *B. fragilis* or control solutions were then gavaged at Day 8 (3 days after DSS cessation). All conventionalized mice were housed in Bioexclusion Racks (Tecniplast). Fecal samples were collected every two weeks or as indicated. Weights were monitored weekly.

### Fecal lipocalin-2 and TNFα measurement

Fecal samples were suspended in PBS containing 1mg/ml protease inhibitor (100mg feces per 1 ml PBS). Sample were thoroughly mixed and supernatant was collected for ELISA.

Mucosal samples were collected by opening intestinal segments along the longitudinal axis and scraping off the mucosal layers with glass slides. Samples were homogenized in 200ul lysis buffer (Cell Signaling Technologies) and [10 mM Tris, pH 7.4, 5 mM MgCl_2_, complete protease inhibitor cocktail (Roche Molecular Biochemicals), 50 U/ml DNase (Amersham), and 50 U/ml RNAse (Ambion)]. Protein concentration for each sample were analyzed with the bicinchoninic acid method.

The protein level of LCN-2 was analyzed using a mouse lipocalin-2/NGAL ELISA kit following the manufacturer’s instructions (R&D Systems). The protein level of TNFα was measured using the Mouse TNFα ELISA kit following the manufacturer’s instructions (ThermoFisher).

### Histological analysis

Cross-section intestinal segments were fixed in 4% formaldehyde. Histological processing and H&E staining were performed by the Human Tissue Resource Center at the University of Chicago. Histological grading of intestinal inflammation was performed by a pathologist blinded to the treatment conditions using a previously validated scoring system as described: no inflammation was scored as 0; modest numbers of infiltrating cells in the lamina propria was scored as 1; infiltration of mononuclear cells leading to separation of crypts and mild mucosal hyperplasia was scored as 2; massive infiltration with inflammatory cells accompanied by disrupted mucosal architecture, loss of goblet cells, and marked mucosal hyperplasia was scored as 3.

### DNA extraction and 16S rRNA gene amplicon sequencing

DNA extraction from feces and colon contents were performed using commercially available kits (Zymo Research). The sequencing of V4 region of the 16S rRNA gene was amplified following the EMP protocol and sequences were obtained by Illumina MiSeq at the Argonne Sequencing Center. The raw sequencing data were analyzed by QIIME2 2019.11 with DADA2^34,35^. Samples with less than 2500 reads were excluded from the analyses. The taxonomy was analyzed using the taxonomy classifier trained on the Greengenes 13_8 99% OTUs that is provided by QIIME2 development team.

For 16S rRNA gene amplicon sequencing analysis, PERMANOVA tests were computed with QIIME2^34^ to assess bacterial composition differences between treatment groups for weighted UniFrac distances data. OTUs between SPF group VS SPF+*B. fragilis* group were compared using Student’s t-test. Statistical significance was achieved at *p* < 0.05.

### 16S whole-length sequencing and taxonomy analysis

Whole length of 16S sequencing were performed with PacBio Single Molecule Real-Time (SMRT) sequencing technology. The service is commercially available through CD-Genomics. 16S whole-length sequences were blasted against 16S rRNA amplicon sequencing data to link the OTUs between the two methods.

### Whole genome sequencing for bacterial DNA and assembling genomes

Bacterial DNA was extracted from a single colony using a microbial DNA extraction kit (QIAGEN, Germantown, MD). Whole genome sequencing for bacterial DNA was performed on Illumina MiSeq at Marine Biological Laboratory (Woods Hole, MA). The raw sequencing data were assembled with PATRIC^36^.

